# Extremely Fast pRF Mapping for Real-Time Applications

**DOI:** 10.1101/2021.03.24.436795

**Authors:** Salil Bhat, Michael Lührs, Rainer Goebel, Mario Senden

**Affiliations:** Department of Cognitive Neuroscience, Faculty of Psychology and Neuroscience, Maastricht University, Maastricht, The Netherlands; Maastricht Brain Imaging Centre, Faculty of Psychology and Neuroscience, Maastricht University, Maastricht, The Netherlands; Department of Research and Development, Brain Innovation B.V., Maastricht, the Netherlands; Department of Neuroimaging and Neuromodeling, Netherlands Institute for Neuroscience, Royal Netherlands Academy of Arts and Sciences (KNAW), Amsterdam, The Netherlands

**Keywords:** population receptive field mapping, real-time fMRI, vision, stimulus encoding

## Abstract

Population receptive field (pRF) mapping is a popular tool in computational neuroimaging that allows for the investigation of receptive field properties, their topography and interrelations in health and disease. Furthermore, the possibility to invert population receptive fields provides a decoding model for constructing stimuli from observed cortical activation patterns. This has been suggested to pave the road towards pRF-based brain-computer interface (BCI) communication systems, which would be able to directly decode internally visualized letters from topographically organized brain activity. A major stumbling block for such an application is, however, that the pRF mapping procedure is computationally heavy and time consuming. To address this, we propose a novel and fast pRF mapping procedure that is suitable for real-time applications. The method is build upon hashed-Gaussian encoding of the stimulus, which significantly reduces computational resources. After the stimulus is encoded, mapping can be performed using either ridge regression for fast offline analyses or gradient descent for real-time applications. We validate our model-agnostic approach *in silico*, as well as on empirical fMRI data obtained from 3T and 7T MRI scanners. Our approach is capable of estimating receptive fields and their parameters for millions of voxels in mere seconds. This method thus facilitates real-time applications of population receptive field mapping.

## 1. Introduction

The retinotopic organization of the human visual cortex has intrigued neuroscientists ever since the beginning of the nineteenth century when visual field maps were first discovered in soldiers suffering from occipital wounds (Fishman, 1997). With the advent of functional magnetic resonance imaging (fMRI) in the early 1990s (Rosen and Savoy, 2012), it became possible to map retinotopy non-invasively (Sereno et al., 1995; DeYoe et al., 1996; Engel et al., 1997). Sereno et al. (1995) pioneered a phase encoding procedure that allowed for the systematic investigation of polar angle and eccentricity distributions. More recently, Dumoulin and Wandell 2008 spearheaded the population receptive field (pRF) mapping approach which provided an expandable, parametric, model of receptive fields. This allowed researchers to study additional properties of receptive fields and their topography as well as relationships between receptive field properties.

The pRF approach has, for instance, enabled researchers to understand the relationship between eccentricity and the size of the receptive fields along the visual hierarchy (Dumoulin and Wandell, 2008; Amano et al., 2009; Harvey and Dumoulin, 2011; Silva et al., 2018), to investigate neural plasticity and visual development from childhood to adulthood (Dekker et al., 2019; Gomez et al., 2018) and to study the dynamic changes of receptive fields in response to attention (de Haas et al., 2014). Furthermore, pRF modelling has aided researchers’ investigations of pathology such as Alzheimer’s disease (Brewer and Barton, 2014), schizophrenia (Anderson et al., 2017), albinism (Ahmadi et al., 2019) and even blindness (Georgy et al., 2019). Additionally, the ability to estimate receptive field parameters is crucial for a number of applications. For instance, receptive fields can serve as a target for transcranial magnetic stimulation (Sack et al., 2009) or provide a spatial forward model for computational models (Peters et al., 2012). Furthermore, receptive fields can be inverted to provide a decoding model for reconstructing perceived, as well as imagined, visual stimuli (Thirion et al., 2006; Senden et al., 2019).

The latter has been suggested to pave the road towards pRF-based brain-computer interface (BCI) communication systems able to directly decode internally visualized letters from topographically organized brain activity (Senden et al., 2019). This is hindered, however, by the method’s immense consumption of computational time and resources. This issue largely remains unaddressed, although some recent work (Thielen et al., 2019) has proposed a fast deep-learning based mapping algorithm (DeepRF). The DeepRF method deploys a deep convolutional neural network (ResNet) which receives a time-series response as input and predicts the corresponding pRF parameters. Once the network is trained, pRF parameters can be estimated simply using a rapid forward pass. This method is indeed faster than standard methods such as grid-search and achieves faithful estimation of pRF parameters with an average computational time of 0.01 to 0.03 seconds per voxel. However, the procedure requires the generated simulated data (for training) and the empirical data to have the same experimental design. Hence, for empirical data with a new experimental design, the network needs to be trained again and the training of the deep neural network can take up to several hours. Moreover, the fMRI data typically contains a large number of voxels. Therefore, despite achieving low computational time per voxel, the total computational time for all voxels is on the order of several minutes. This makes the approach unfeasible for real-time analysis. With the aim to enable estimation of receptive fields in real-time, we propose here a novel model-agnostic procedure which can be used offline (using ridge regression) as well as online (using gradient descent).

The method relies on regularized linear regression whose basis set is a hashed-Gaussian encoding of the stimulus-evoked response. Specifically, the stimulus space is exhaustively partitioned as a set of features where each feature uniquely encodes the stimulus by computing the overlap between the stimulus and a set of randomly positioned Gaussians. This type of encoding considerably reduces the memory requirements with a low performance loss and thereby accelerates the calculations.

Using two previously acquired datasets from 3 Tesla and 7 Tesla MR systems, we show that the proposed approach works extremely fast. It is able to estimate receptive field shapes of millions of voxels within seconds. This allows the selection of visually responsive voxels through cross-validation and subsequent estimation of receptive field parameters within about one minute even if the data consists of more than 4 million voxels.

## 2. Methods

### 2.1. Fast Mapping Procedure

#### Tile Coding and Hashing

To reduce computation time as well as to lower memory requirements, we encode the stimulus using tile coding and hashing (Albus, 1975, 1981). Tile coding is a linear function approximation used in reinforcement learning (Sutton et al., 1998) to deal with large and continuous state spaces. In tile coding, the state space is exhaustively partitioned into subregions called tiles. Usually, the presence of an entity within a tile (in this case, the presence of a stimulus in a region of the visual field) is encoded in a binary fashion. However, it is also possible to encode features using radial basis functions which have the additional benefit of varying smoothly. Memory requirements can be reduced further by hashing a group of individual, non-contiguous, tiles into a single tile. Figure 1 depicts tile-coding and hashing of sample stimuli. The presence of a stimulus is encoded as the extent of overlap between the stimulus and hashed tilings. For our purposes, we use a 2-D isotropic Gaussian as the radial basis function. Subsequently, we hash by combining five randomly selected Gaussians into a single tile leading to a total of 250 tilings. The 5 Gaussian tiles within a tiling may or may not overlap. We normalized each tile to ensure that the area under its surface is equal to one. The code used in this paper is publically available at https://github.com/ccnmaastricht/real_time_pRF

**Figure 1:**
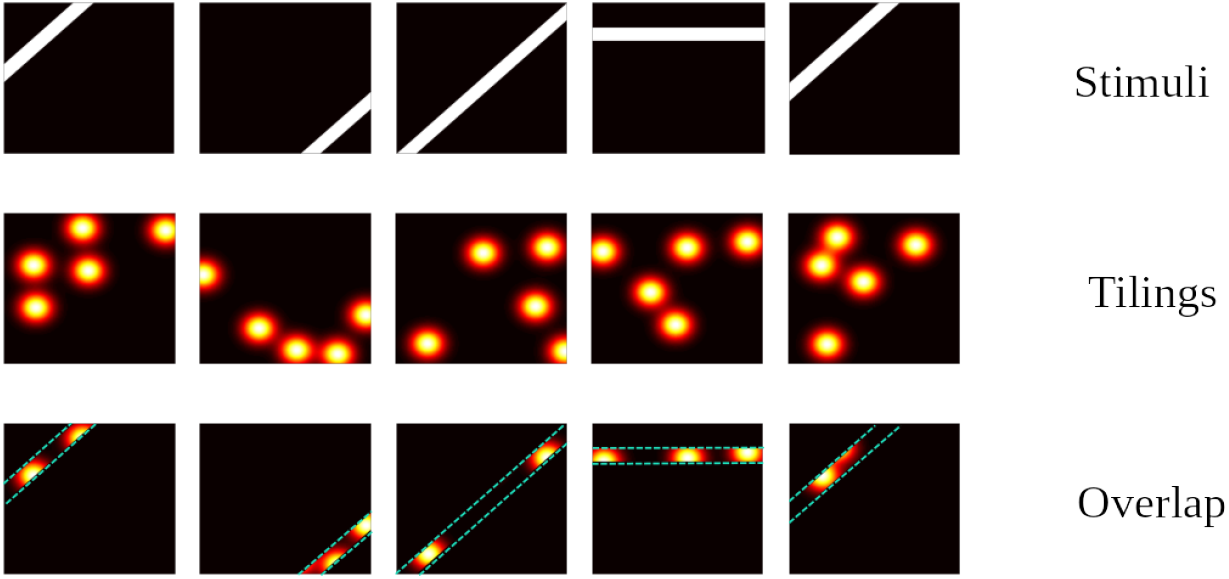
Illustration of tile-coding and hashing. The top rows shows sample stimuli. The middle row shows sample tilings, each corresponding to a stimulus and each containing 5 Gaussians which make up one tile. The bottom row shows overlap between stimuli and corresponding tiles.

#### Encoding Stimuli

Using hashed-Gaussians as tiles, it is possible to encode retinotopic stimuli. First, an overlap between a binary indicator function and a tiling matrix Γ(pixels-by-tiles) is computed. The binary indicator function marks the position of the stimulus aperture at each moment in time **S** (time-by-pixels). Subsequently, the computed overlap is convolved with a canonical two-gamma hemodynamic response function (HRF) function (*h*) to obtain the encoded stimulus *ϕ*

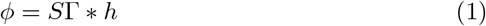

#### Ridge Regression

We use ridge regression for fast offline pRF mapping (i.e. after all functional volumes have been acquired). Specifically, the BOLD activity response is modeled by

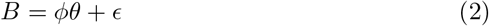

where *θ* are the estimated weights and *ϵ* denotes the residuals. Note that, prior to computing *θ*, both *ϕ* and the BOLD data *B* are z-normalized. In order to estimate *θ*, the discrepancy between the measured and predicted BOLD response (*ϕθ*) needs to be minimized. Therefore, we define the error or the loss as

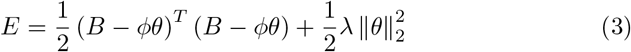

In order to avoid over-fitting, we use *L*_2_ regularization and *λ* denotes the regularization factor. The gradient of the error with respect to *θ* can be computed as

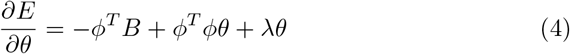

By setting 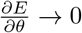 and solving for optimal *θ*, we get

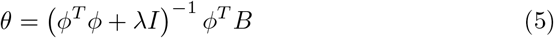

Receptive fields can now be straightforwardly obtained by multiplying the tiling matrix with the estimated *θ*: *W* = Γ*θ*. These *raw* receptive fields are then subjected to post-processing. The raw receptive fields contain anomalous pixel intensities. These can be removed by first normalizing the raw receptive fields to the range [0, 1] and then shrunk by raising them to a power of some positive integer (shrinkage factor). This shrinks noisy pixel intensities close to 0 while leaving those close to 1 unaffected (figure 2), thus yielding cleaner receptive fields.

**Figure 2:**
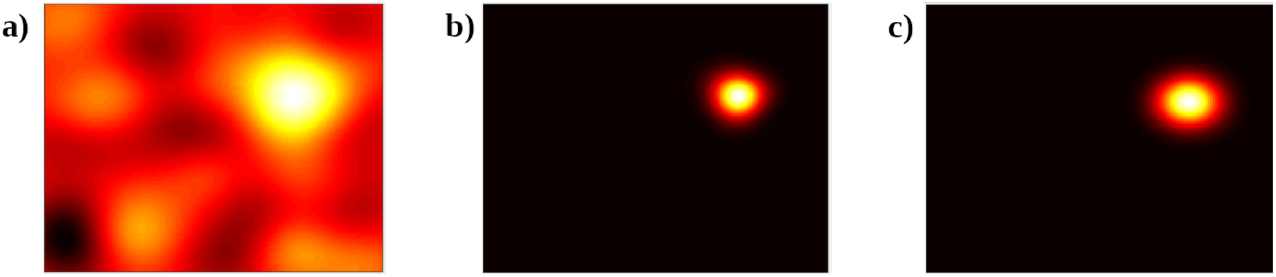
The effect of shrinking a raw receptive field. **a**, Raw receptive field displaying undesirably large pixel intensities. **b**, The receptive field after shrinkage with a factor of 9. **c**, The corresponding ground truth receptive field.

#### 2.1.1. Similarity Metric

In order to compare the receptive fields obtained from ridge regression with corresponding ground-truth/grid-search receptive fields, we use the Jaccard Index (or Jaccard Similarity). Since the Jaccard Index (JI) is a conservative metric, we derive a *Null-model* from a resampling procedure for a better interpretation. Specifically, for each estimated receptive field, we pair it with a random ground-truth/grid-search receptive field and compute the JI. The average over these pairs is the JI of one randomization. We repeat this procedure 1000 times to obtain a Null-distribution of randomized JIs. We refer to the mean of Null-distribution as the *baseline.*

### 2.2. Online Gradient Descent

For online pRF mapping we use gradient descent to iteratively update *θ* with each acquired volume. In this case, we define the loss function as

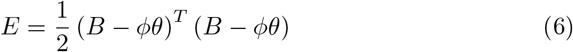

The gradient of the loss function with respect to the parameter *θ* is

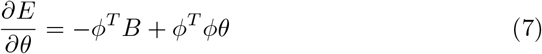

At each time point, *θ* is updated by a factor (learning rate *η*) of the gradient. Note that, unlike ridge regression, a regularization term is not needed in this case, as gradient descent is effectively regularized by the learning rate (see Appendix A). Considering the *n*^*th*^ time point, the update can be computed as

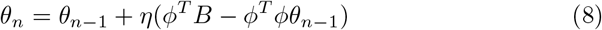

Similar to the offline method, prior to tile coding and hashing, the stimulus needs to be convolved with the HRF. Furthermore, both the BOLD signal *B* and encoded stimulus *ϕ* need to be z-normalized. However, in an online setting this needs to be performed in real-time. Real-time z-normalization requires realtime estimation of the mean and variance of a signal which can be done using Welford’s online algorithm (Welford, 1962). Once the current mean 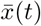 and variance *σ*^2^(*t*) have been estimated, the current z-score can be estimated as

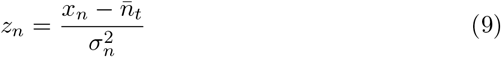

#### Voxel Selection

Since not all measured voxels are visual, and hence may not carry significant information, a voxel selection procedure is desirable. We evaluate voxels in terms of the cross-validated Pearson correlation coefficient (fitness) between their predicted and measured BOLD responses. To account for temporal auto-correlation in the BOLD response, we use a blocked cross-validation procedure (Roberts et al., 2017). Specifically, the data is split into *p* windows along the time axis. Ridge regression is performed on window 1 and the estimated *θ* values are used to predict the BOLD response for the remaining *p* − 1 windows. This is followed by ridge regression on windows 1 and 2 and predicting the BOLD response in the remaining *p* − 2 windows. This procedure continues until ridge regression is performed on windows 1 to *p* − 1 and the BOLD response is predicted for the *p*_*th*_ (last) window. The overall fitness for each voxel is then given by the mean of fitness values computed for each split. The data used in this paper has 304 time points. We split the data into 4 windows of equal length and retain voxels whose fitness falls within the top 1 %.

### 2.3. Fast pRF Parameter Estimation

Post-processed receptive fields obtained from our ridge regression and gradient descent methods can be readily used to estimate parameters of an isotropic Gaussian pRF model (i.e. the x-location, y-location and size) using a fast procedure. Since peak pixel intensity of a Gaussian receptive field is at its center, we estimate the x- and y-coordinate of pre-processed model-free receptive fields by finding the location of their peak pixel intensity. To estimate the size of receptive fields, our procedure utilizes the relationship between the standard deviation, eccentricity and the mean pixel intensity in an isotropic Gaussian embedded in a finite image. Specifically, given a Gaussian at a fixed location, mean pixel intensity increases as a function of its standard deviation. Furthermore, in a finite image and assuming a fixed size, mean pixel intensity decreases as the Gaussian is progressively moved toward the edge of an image. Therefore, for a given image size, we generate isotropic Gaussians with 25 different standard deviations, located at 25 eccentricities along an axis of 45°, and compute their mean pixel intensities. This can be utilized to perform a linear regression with mean pixel intensity and eccentricities predicting the receptive field size. We then use the resulting regression weights together with previously estimated locations and mean pixel intensity of our receptive fields to obtain an estimate of their size.

### 2.4. Data

#### 2.4.1. Simulated Data

We simulate fMRI data for a V1-like cortical sheet extending 55 mm along and approximately 40 mm orthogonal to the horizontal meridian in both hemispheres. Since such a sheet is akin to a flattened cortical mesh, model units are referred to as vertices rather than voxels. Each vertex in the model is a 0.5 mm isotropic patch whose receptive field center is directly related to its position on the surface in accordance with a complex-logarithmic topographic mapping (Schwartz, 1980; Balasubramanian et al., 2002) with parameter values (*a* = 0.7, *α* = 0.9; Polimeni et al., 2005). The shape of model receptive fields is given by a 2-dimensional Gaussian

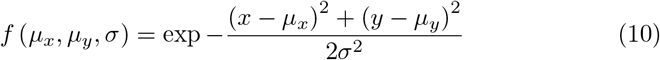

with (*μ*_*x*_, *μ*_*y*_) being the receptive field center and *σ* its size. Below an eccentricity of *e* = 2.38 all model vertices have a receptive field size of *σ* = 0.5 whereas they exhibit a linear relationship with eccentricity (*σ* = 0.21*e*) beyond this cutoff (c.f. Freeman and Simoncelli, 2011).

A simulated fMRI signal (sampled at a rate of 0.5 Hz) for each vertex is obtained by first performing element-wise multiplication between the receptive field of a vertex and the effective stimulus presented per time point, summing the result and subsequently convolving the obtained signal with the canonical two-gamma hemodynamic response function. Two sources of distortion are added to the signal. First, a spatial smoothing kernel is applied to simulate the point-spread function of BOLD activity on the surface of the striate cortex (Shmuel et al., 2007). Second, autocorrelated noise generated by an Ornstein-Uhlenbeck process with variance 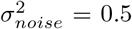 is added. The smoothing kernel is independently applied to the clean signal and the noise before the two are combined. We simulate both 3T- and 7T-like signals by adjusting the full-width at half-maximum of the spatial smoothing kernel (3.5 mm and 2 mm for 3T and 7T, respectively; c.f. Shmuel et al., 2007) and the time constant of the Ornstein-Uhlenbeck process (2.25 s and 1 s for 3T and 7T, respectively).

#### 2.4.2. Three Tesla Empirical Data

This dataset, previously described in (Senden et al., 2014), comprises a retinotopy run obtained from three participants (all male, age range = 27-35 years, mean age = 32 years). During this run a bar aperture (1.5° wide) revealing a flickering checkerboard pattern (10 Hz) was presented in four orientations. For each orientation, the bar covered the entire screen in 12 discrete steps (each step lasting 2 s). Within each orientation, the sequence of steps (and hence of the locations) was randomized and each orientation was presented six times. Furthermore, within each presentation four bar stimuli were replaced with mean luminance images for four consecutive steps. These data were acquired on a Siemens 3T Tim Trio scanner equipped with a 32-channel head coil (Siemens, Erlangen, Germany) using a gradient-echo echo-planar imaging sequence (31 transversal slices; TR = 2000 ms; TE = 30 ms; FA = 77°; FoV = 216 x 216 mm^2^; 2 mm isotropic resolution; no slice gap; GRAPPA = 2) and are publicly available (Senden et al., 2014). Preprocessing consisted of slice scan time correction, (rigid body) motion correction, linear trend removal, and temporal high-pass filtering (up to 2 cycles per run).

#### 2.4.3. Seven Tesla Empirical Data

This dataset, previously described in (Senden et al., 2019), comprises retinotopy as well as passive viewing of letter stimuli obtained from six participants (2 female, age range = 21-49 years, mean age = 30.7 years). During the retinotopy run a bar aperture (1.33° wide) revealing a flickering checkerboard pattern (10 Hz) was presented in four orientations. For each orientation, the bar covered the entire screen in 12 discrete steps (each step lasting 3 s). Within each orientation, the sequence of steps (and hence of the locations) was randomized and each orientation was presented six times. During the passive viewing run four letters (‘H’, T’, ‘S’ and ‘C’) were presented in a 8° by 8° bounding frame for a duration of 6 s and their shape was filled with a flickering checkerboard pattern (10 Hz). These data were acquired on a Siemens Magnetom 7T scanner (Siemens; Erlangen, Germany) equipped with a 32 channel head-coil (Nova Medical Inc.; Wilmington, MA, USA) using high-resolution gradient echo echoplanar imaging sequence (82 transversal slices; TR = 3000 ms; TE = 26 ms; generalized auto-calibrating partially parallel acquisitions (GRAPPA) factor = 3; multi-band factor = 2; FA = 55; FoV = 186 x 186 mm^2^; 0.8 mm isotropic resolution). In addition, this dataset includes five functional volumes acquired with opposed phase encoding directions to correct for EPI distortions that occur at higher field strengths (Andersson et al., 2003). Preprocessing further consisted of (rigid body) motion correction, linear trend removal, and temporal high-pass filtering (up to 3 cycles per run).

For visualization purposes, we also include anatomical data for subject 3. Anatomical data was acquired with a T1-weighted magnetization prepared rapid acquisition gradient echo (Marques et al., 2010) sequence [240 sagittal slices, matrix = 320 x 320 m, voxel size = 0.7 mm isotropic, first inversion time TI1 = 900 ms, second inversion time TI2 = 2750 ms, echo time (TE) = 2.46 ms repetition time (TR) = 5000 ms, first nominal flip angle = 5°, and second nominal flip angle = 3°. Anatomical images were interpolated to a nominal resolution of 0.8 mm isotropic to match the resolution of functional images. In the anatomical images, the grey/white matter boundary was detected and segmented using the advanced automatic segmentation tools of BrainVoyager 20 which are optimized for high-field MRI data. A region-growing approach analyzed local intensity histograms, corrected topological errors of the segmented grey/white matter border, and finally reconstructed meshes of the cortical surfaces (Kriegeskorte and Goebel, 2001; Goebel et al., 2006)

### 2.5. Real-time Processing

To mimic a real-time scenario, we limited the preprocessing to trilinear 3D rigid body motion correction which was applied in a simulated real-time setup using Turbo-BrainVoyager (TBV) (v4.0b1, Brain Innovation B.V., Maastricht, The Netherlands). The data was accessed directly from TBV using a network interface providing fast transfer speed suitable for real-time applications. The receiver was implemented in MATLAB™(version 2019a, The Mathworks .inc, Natick, MA, USA) using JAVA based TCP/IP interfaces.

## 3. Results

All experiments were performed using MATLAB™(version 2019a, The Mathworks .inc, Natick, MA, USA) running on an HP^®^ Z440 workstation with an Intel^®^ Xeon^®^ Processor (E5-1650 v4, 32GB RAM) and an Ubuntu 20.04 operating system. The set of hyperparameters (learning rate *η* = 0.1, shrinkage factor = 6 and *FWHM* = 0.15) remain the same for all experiments, except for reconstruction of perceived letter shapes where a shrinkage factor of 9 was used. All the figures generated using MATLAB™(including the parts of Figures 1 and 2) were generated using *export_fig* (Altman, 2020).

### 3.1. Fast Mapping Procedure

#### 3.3.1. Simulated Data

The fast, ridge-based, mapping procedure was first tested on simulated data to investigate whether it faithfully recovers known population receptive field shapes and their parameters. Overall, the mean Jaccard Similarity (JS) between the estimated and ground-truth receptive field shapes was 0.3452 (95 % CI [0.3409, 0.3495]) and 0.3920 (95 % CI [0.3877, 0.3963]), for simulated 3T and 7T data, respectively. For comparison, corresponding Null-model JS values were 0.0418 and 0.0410, respectively. There is thus good correspondence between estimated and ground-truth receptive field shapes which is also apparent from the sample receptive fields shown in figure 3. Next, we examined the correspondence between receptive field parameters obtained with the two methods. While receptive fields mapped using the ridge regression are not exactly Gaussian, estimated parameters nevertheless show an excellent correspondence with ground truth parameters for both simulated 3T and 7T data (see figures 4 and C.18 respectively as well as table 1). Please note that despite the high correlation, the receptive field size tends to be slightly overestimated by our method. The size of mapped receptive fields can be adjusted using the shrinkage factor. However, for the sake of comparison, we use a constant shrinkage factor across the datasets.

**Table 1:** Correlations between estimated and ground-truth pRF parameters

| | X-coordinate | Y-coordinate | size ($\sigma$) |
| --- | --- | --- | --- |
| 3T | 0.9913 (95 % CI [0.9909, 0.9916]) | 0.9871 (95 % CI [0.9865, 0.9876]) | 0.9674 (95 % CI [0.9660, 0.9686]) |
| 7T | 0.9958 (95 % CI [0.9957, 0.9960]) | 0.9949 (95 % CI [0.9946, 0.9951]) | 0.9681 (95 % CI [0.9668, 0.9693]) |

**Table 2:**
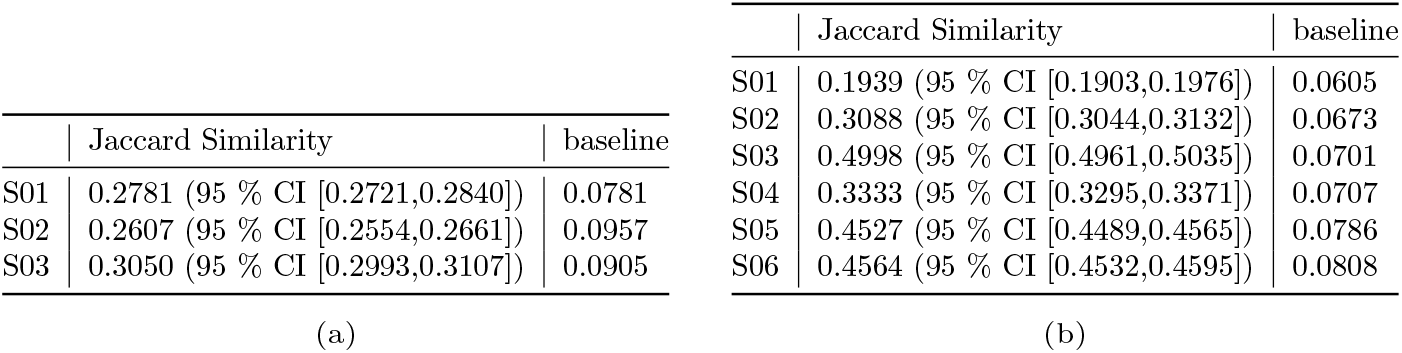
Mean Jaccard Similarities between receptive fields estimated with the fast procedure and those obtained from grid-search for **a)** 3T and **b)** 7T empiricial data.

**Figure 3:**
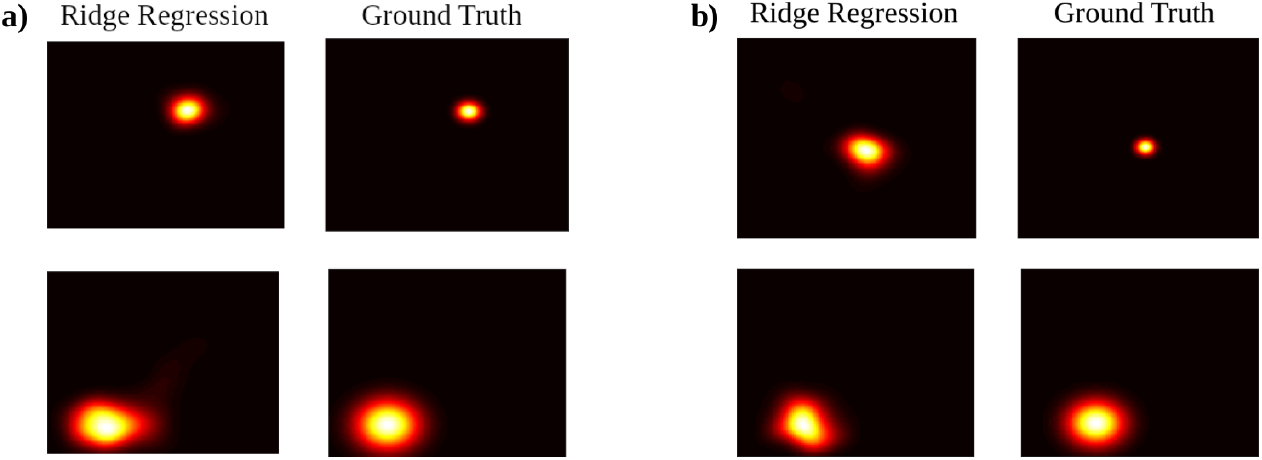
Comparison of ridge-estimated and ground-truth receptive fields. **a)** Small (top) and large (bottom) estimated and ground-truth receptive fields for simulated 3T data (*TR* = 2000*ms*). **b)** Small (top) and large (bottom) estimated and ground-truth receptive fields for simulated 7T data (*TR* = 3000*ms*).

**Figure 4:**
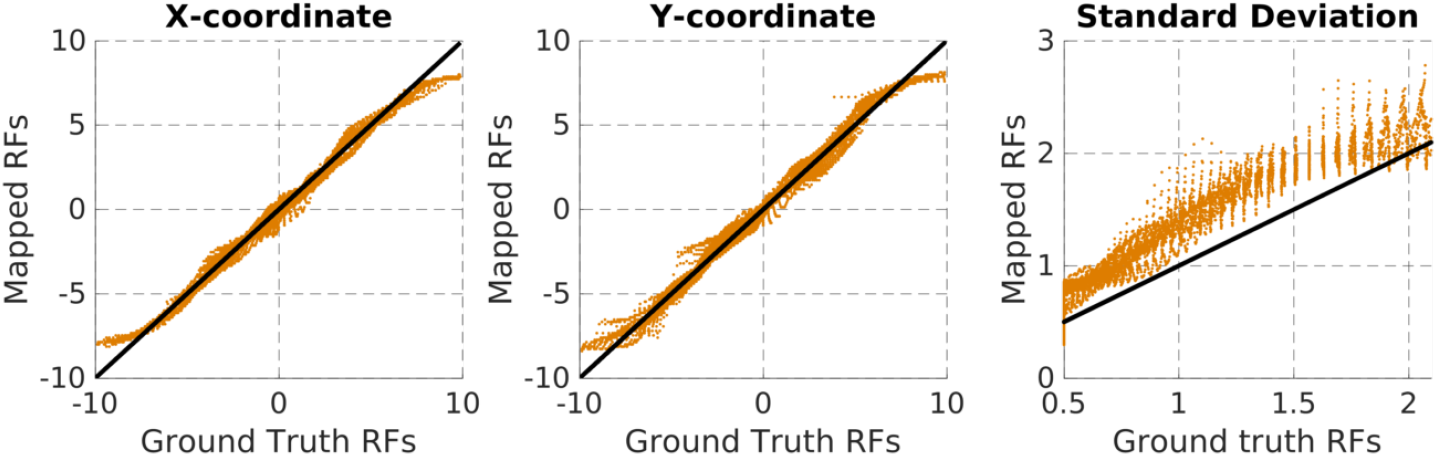
Estimated vs ground-truth pRF parameters. A line with a slope of 1 is included as a reference. Voxels whose receptive fields lie outside the field of view were ignored for estimating pRF parameters. Results are from simulated 3T data. Results for simulated 7T data are comparable (see supplementary figure C.18).

Next, we evaluated the ridge-based mapping approach in terms of its computational performance. To that end we measured both memory consumption and the computational time required for the mapping procedure itself as well as for subsequent parameter estimation. Computational times were estimated using MATLAB™’s stopwatch utility. The execution time measured using this utility can be affected by many unknown variables pertaining to memory, processor, caching in memory, MATLAB™’s just-in-time compiler, etc. This may influence the execution time measurement each time a subroutine is executed. Therefore, we report computational times as a mean over 100 runs. Memory requirements were estimated using GNU/Linux’s *pmap* command. The memory requirements reported here are calculated as *mem*_*max*_ − *mem*_0_, where *mem*_*max*_ is the maximum amount of memory consumed during the procedure and *mem*_0_ is the memory occupied by MATLAB™before starting the procedure (which includes loading of data into memory and other background processes occupying memory). Memory consumption during the procedure was logged every 0.1 seconds using GNU/Linux’s *watch* command. Note that since here we are only interested in computational performance we test the mapping procedure on randomly generated data of the size 304 *by voxels*. Memory consumption was averaged over 100 repetitions of the procedure. As can be appreciated from figure 5 the ridge-based mapping procedure is extremely fast (less than 10 s). The computational time only starts to increase as the needed memory exceeds the available memory. As a consequence, virtual memory gets consumed which slows down the mapping procedure. Memory consumption scales linearly with the number of voxels and allows for estimation of ~ 1.75 and ~ 3.5 million voxels on systems with 8GB and 16GB of RAM, respectively.

**Figure 5:**
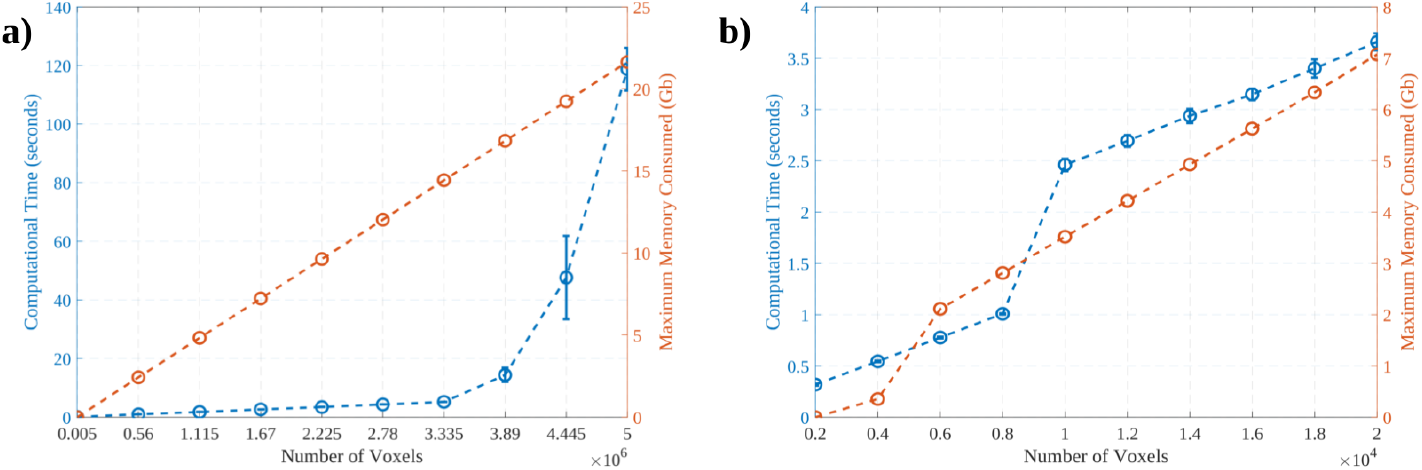
Memory consumption (orange) and computational time (blue), as a function of the number of voxels for **a)** ridge regression and **b)** parameter estimation. Data points corresponding to 0s reflect < 1 Kb.

#### 3.1.2. Empirical Data

Following up on simulation results, we tested the ridge-based mapping procedure on previously acquired empirical data. Similar to the simulated data, we asses our method in terms of its ability to estimate pRF shapes and their parameters as well as computational performance. Since ground truth receptive field shapes and parameters are not known for empirical data, we assess our method on its ability to produce estimates that are consistent with a grid-search pRF mapping procedure. Sample receptive fields estimated in the 3T and the 7T empirical data are shown in figures 6 and 7, respectively. Retinotopic surface maps for a representative subject in the 7T dataset are shown in figure 8. These results qualitatively indicate a good agreement between receptive field shapes and parameters between our method and the grid-search approach. Quantitatively, we observe that the Jaccard similarity between receptive fields estimated using the ridge-based and grid-search methods consistently exceed those expected based on the Null model. The JS is particularly high for subjects 03, 05 and 06 for the 7T empirical dataset. In terms of correspondence between the pRF parameters obtained from the fast procedure and those obtained from grid-search, the correlation coefficients shown in Table 3 indicate that correspondence is generally good. This is also apparent from scatter plots showing the correspondence between pRF parameters obtained from our method and grid-search in representative subjects (see 9**a** and 9**b** for the 3T and 7T dataset, respectively).

**Table 3:** Correlation coefficients between the pRF parameters obtained from the fast procedure and those obtained from grid-search for **a)** 3T and **b)** 7T datasets, respectively.

|  | X-coordinate | Y-coordinate | Standard Deviation |
| --- | --- | --- | --- |
| S01 | 0.7557 (95 % CI [0.7436, 0.7674]) | 0.7381 (95 % CI [0.7252, 0.7505]) | 0.2287 (95 % CI [0.2023, 0.2548]) |
| S02 | 0.6906 (95 % CI [0.6758, 0.7048]) | 0.7526 (95 % CI [0.7404, 0.7644]) | 0.2809 (95 % CI [0.2552, 0.3062]) |
| S03 | 0.7551 (95 % CI [0.7429, 0.7667]) | 0.7270 (95 % CI [0.7137, 0.7398]) | 0.1758 (95 % CI [0.1488, 0.2026]) |

(a)
|  | X-coordinate | Y-coordinate | Standard Deviation |
| --- | --- | --- | --- |
| S01 | 0.6411 (95 % CI [0.6294, 0.6525]) | 0.6042 (95 % CI [0.5916, 0.6165]) | 0.0386 (95 % CI [0.0190, 0.0581]) |
| S02 | 0.7038 (95 % CI [0.6937, 0.7135]) | 0.6238 (95 % CI [0.6116, 0.6356]) | -0.0298 (95 % CI [-0.0494, -0.0102]) |
| S03 | 0.9723 (95 % CI [0.9712, 0.9734]) | 0.9645 (95 % CI [0.9632, 0.9659]) | 0.5693 (95 % CI [0.5559, 0.5824]) |
| S04 | 0.8332 (95 % CI [0.8271, 0.8391]) | 0.7761 (95 % CI [0.7682, 0.7838]) | 0.0970 (95 % CI [0.0776, 0.1164]) |
| S05 | 0.9066 (95 % CI [0.9030, 0.9100]) | 0.8976 (95 % CI [0.8937, 0.9013]) | 0.3332 (95 % CI [0.3157, 0.3505]) |
| S06 | 0.9216 (95 % CI [0.9186, 0.9245]) | 0.9249 (95 % CI [0.9220, 0.9277]) | 0.3229 (95 % CI [0.2941, 0.3295]) |

**Figure 6:**
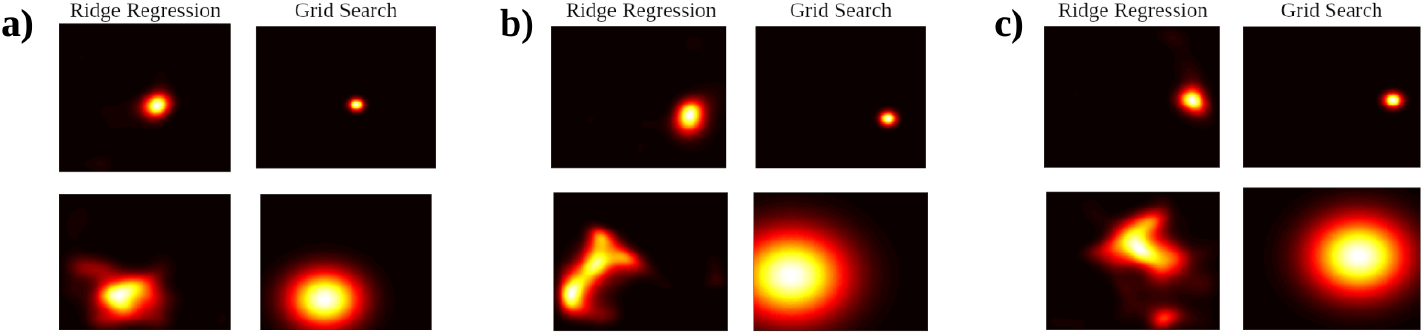
Comparison of ridge-estimated and ground-truth receptive field parameters for 3T data. **a)** Small (top) and large (bottom) estimated and ground-truth receptive fields for subject 1. **b,c)** Same as panel **a** for subjects 2 and 3, respectively

**Figure 7:**
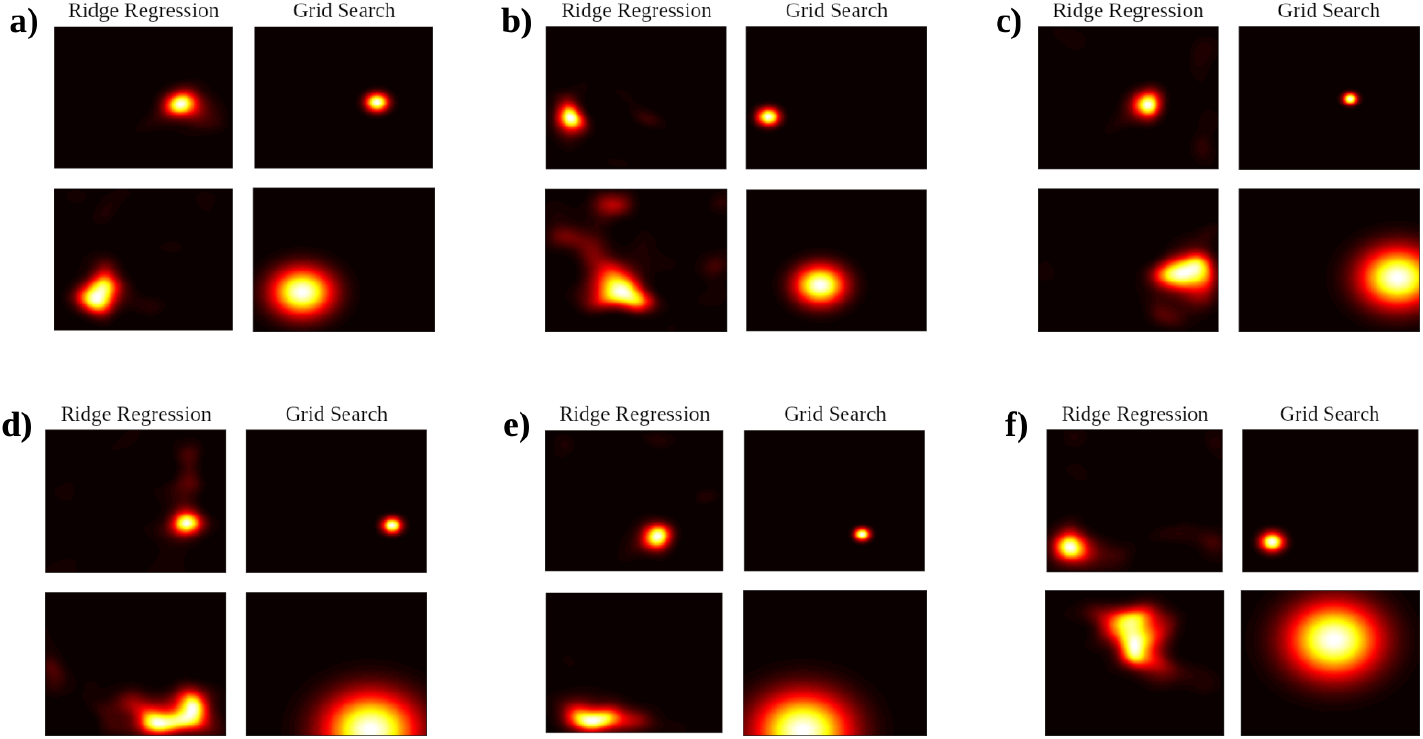
Comparison of ridge-estimated and ground-truth receptive field parameters for 7T data. **a)** Small (top) and large (bottom) estimated and ground-truth receptive fields for subject 1. **b-f)** Same as panel **a** for subjects 2 to 6, respectively

**Figure 8:**
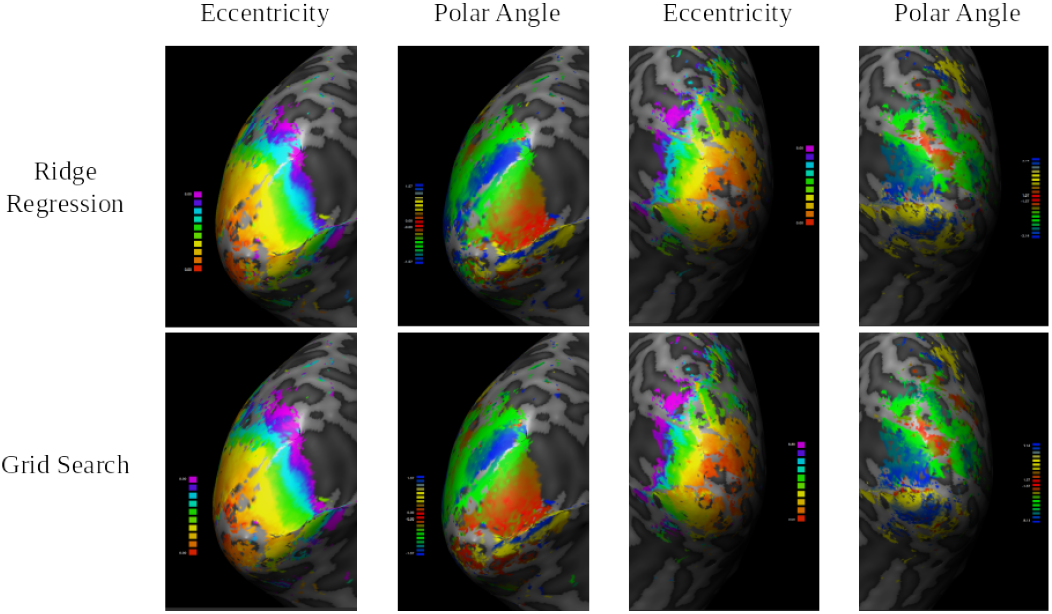
Exemplary eccentricity and polar angle maps in both hemispheres of subject 3 in the 7T dataset. The upper row shows maps obtained using our fast parameter estimation procedure whereas the bottom row shows maps obtained using a grid-search procedure. In accordance with the correlation results between maps (see table 3b), the two polar angle and eccentricity maps are visually highly similar.

**Figure 9:**
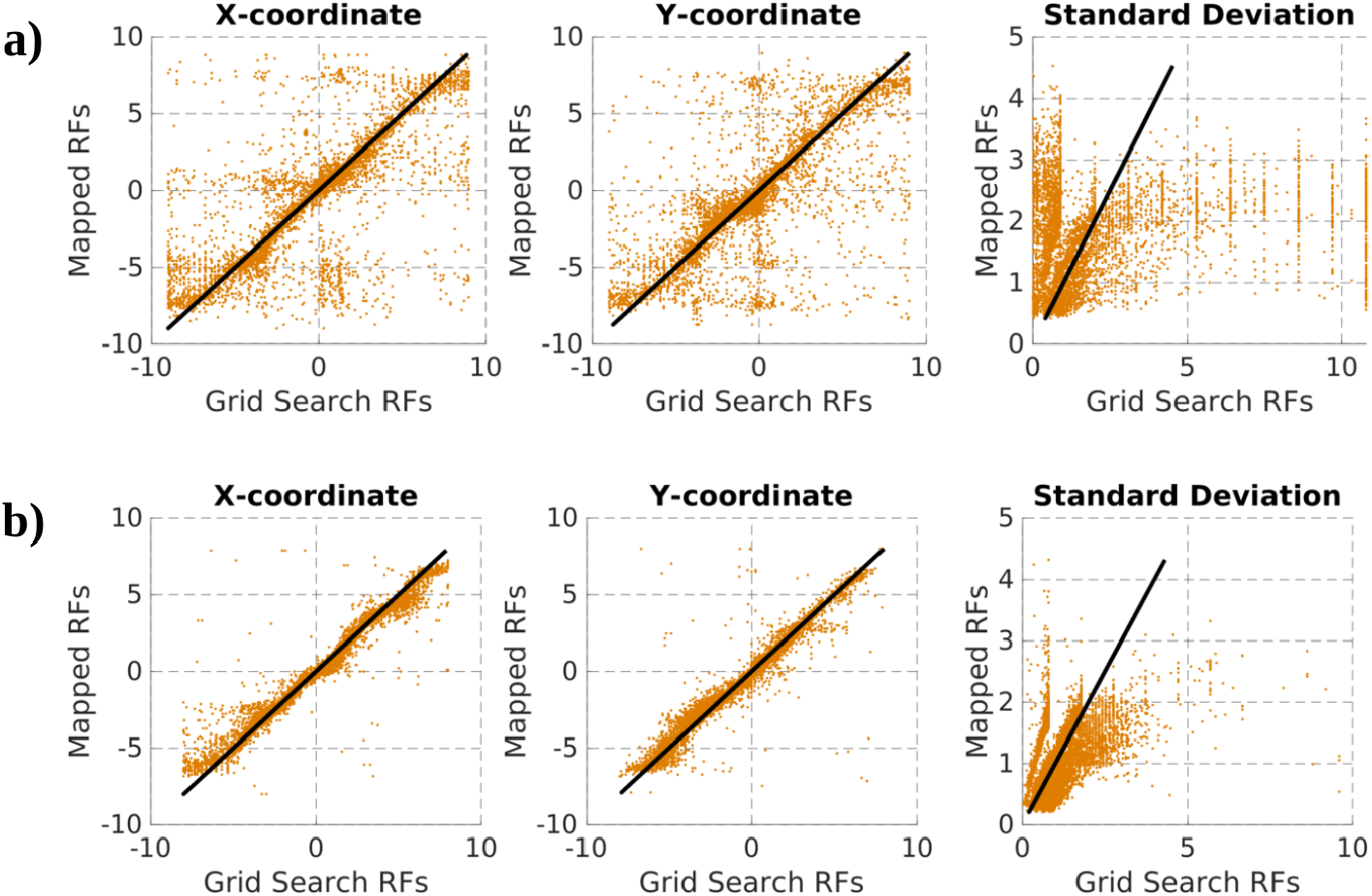
Fast procedure vs grid-search estimated pRF parameters for **a)** 3T (subject 1) and **b)** 7T (subject 3) data, respectively. A line with a slope of 1 is included as a reference.

We again evaluate computational performance in terms of computational times and memory consumption. We estimate both based on 100 runs for each dataset. Since each subject has a different number of voxels, for each run a subject was chosen randomly. The computational time is computed separately for ridge regression, cross-validation and parameter estimation. The mean computational times or execution times for both datasets are reported in tables 4a and 4b.

**Table 4:**
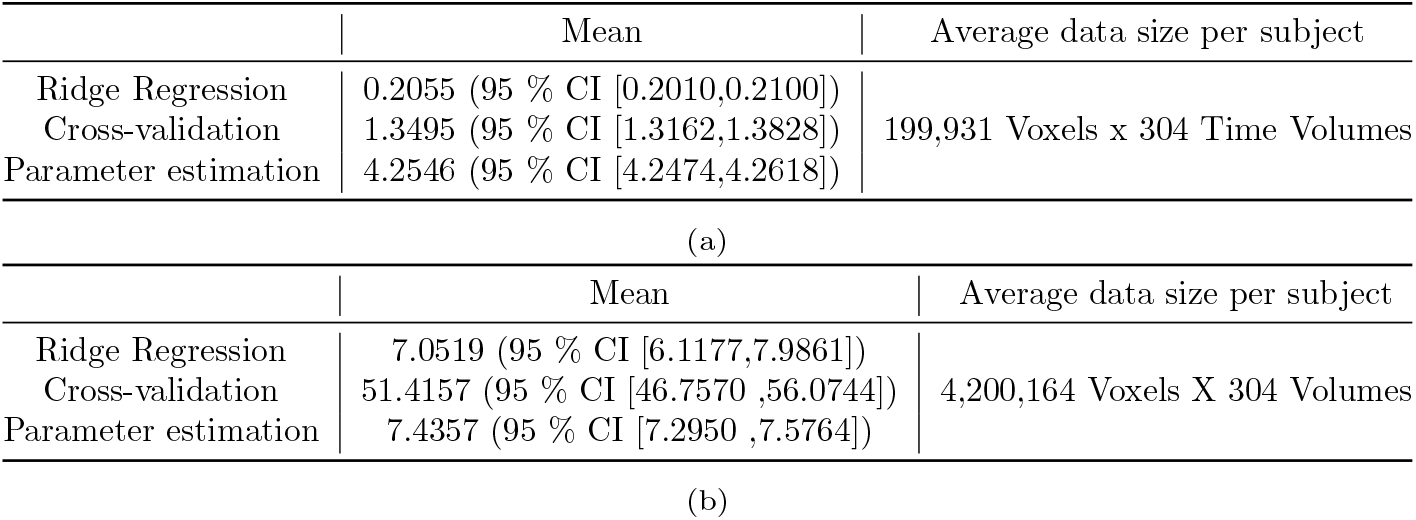
Mean computational times in seconds for **a)** 3T and **b)** 7T empirical data. The average data size per subject reflects the amount of data processed by the algorithm at a time.

The computational times suggest that our algorithm is extremely fast in mapping receptive fields. The actual mapping procedure happens within a second for the 3T dataset and in a few seconds for the 7T dataset. Cross-validation, which selects the best voxels, finishes in a couple of seconds for the 3T dataset and takes less than a minute for the 7T dataset. The estimation of pRF parameters (for all voxels) also takes only a few seconds for both datasets. This means that receptive fields and their pRF parameters are readily available for further analysis.

### 3.2. Online Gradient Descent

To demonstrate the capability of online gradient descent to work in a real-time setting, we mimicked a real-time scenario using TurboBrainVoyager™(as described in section 2.5). We show in Appendix A that ridge regression and online gradient descent yield similar receptive fields through hyperparameter sharing. Therefore, we do not provide an evaluation of the ability of the method to reliably estimate receptive field shapes and parameters. Instead, we evaluate its performance in terms of whether estimated receptive fields are suitable for projecting cortical activity back into the visual field. For that purpose we utilize data acquired as subjects passively viewed letter shapes previously described in (Senden et al., 2019). The reconstructions obtained from our approach (see figure 10) are recognizable and comparable those obtained from receptive fields resulting from grid-search.

**Figure 10:**
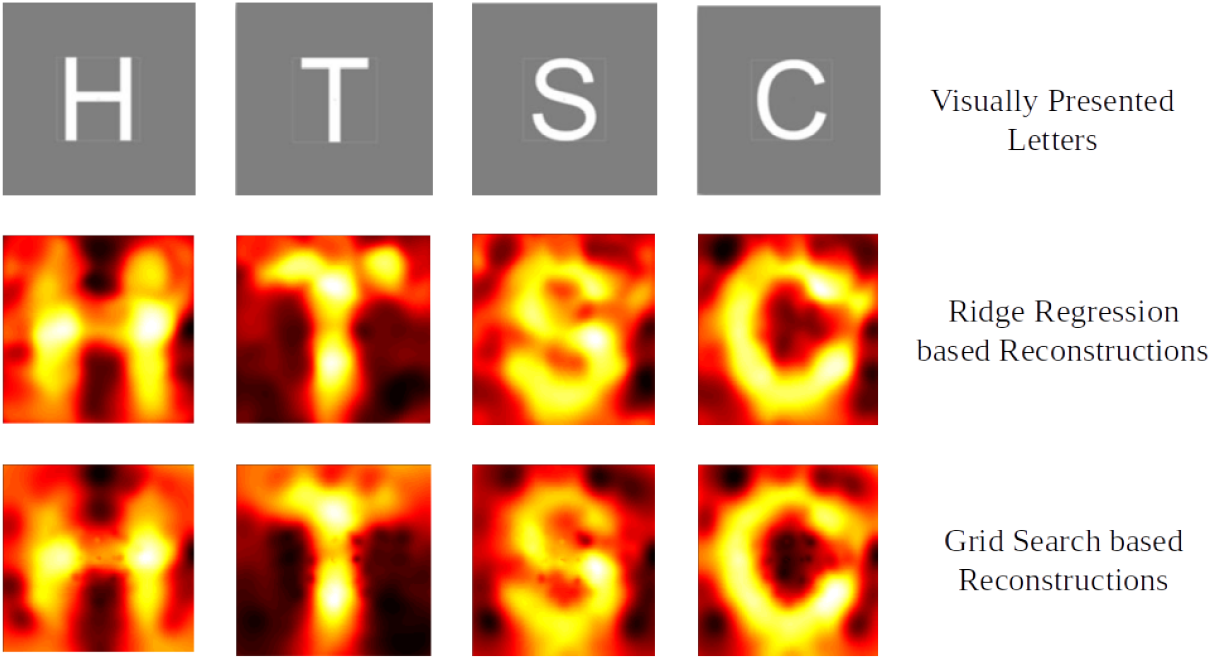
Reconstructions of perceived letter shapes.

The mean computational times per time volume per subject for the 3T and 7T datasets are reported in tables 5a and 5b, respectively. MATLAB™uses a just-in-time compiler, which has to be executed the first time and has to first load the subroutine into memory and compile. This often causes the first iteration to be slower. Therefore, we exclude the execution time of the first time volume while computing the mean and standard deviation and report it separately. The average computational time per acquired volume is less than the repetition time (2000*ms* for 3T and 3000*ms* for 7T), which means that the receptive fields are updated before the next time volume is acquired. This is especially useful in a real-time setting where analysis needs to be done as the data is being acquired. Figure 11 depicts how memory requirements scale with computational time. The computational only starts to increase when needed memory exceeds the available memory. Generally, up to 1 million voxels can be comfortably estimated within less than 1500*ms* and requiring less than 2GB of RAM.

**Table 5:**
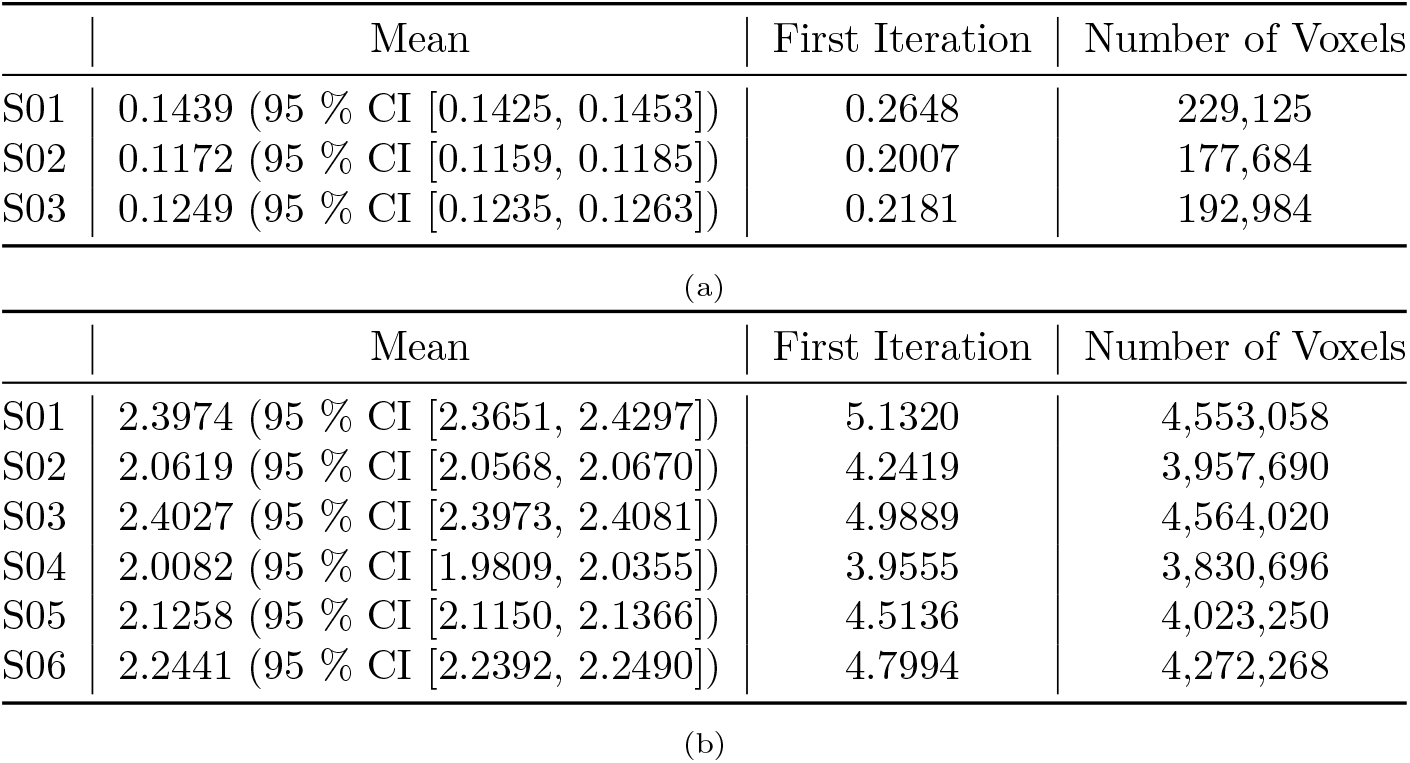
Mean computational time (in seconds) per time volume per subject for the real-time mapping technique performed on **a)** 3T and **b)** 7T empirical data. For each subject, data for 304 time volumes was recorded. Since the first iteration (corresponding to the first time volume) is usually abnormally high, it is not included for computing mean and standard deviation.

|  | Mean | First Iteration | Number of Voxels |
| --- | --- | --- | --- |
| S01 | 0.1439 (95 % CI [0.1425, 0.1453]) | 0.2648 | 229,125 |
| S02 | 0.1172 (95 % CI [0.1159, 0.1185]) | 0.2007 | 177,684 |
| S03 | 0.1249 (95 % CI [0.1235, 0.1263]) | 0.2181 | 192,984 |

(a)
|  | Mean | First Iteration | Number of Voxels |
| --- | --- | --- | --- |
| S01 | 2.3974 (95 % CI [2.3651, 2.4297]) | 5.1320 | 4,553,058 |
| S02 | 2.0619 (95 % CI [2.0568, 2.0670]) | 4.2419 | 3,957,690 |
| S03 | 2.4027 (95 % CI [2.3973, 2.4081]) | 4.9889 | 4,564,020 |
| S04 | 2.0082 (95 % CI [1.9809, 2.0355]) | 3.9555 | 3,830,696 |
| S05 | 2.1258 (95 % CI [2.1150, 2.1366]) | 4.5136 | 4,023,250 |
| S06 | 2.2441 (95 % CI [2.2392, 2.2490]) | 4.7994 | 4,272,268 |

**Figure 11:**
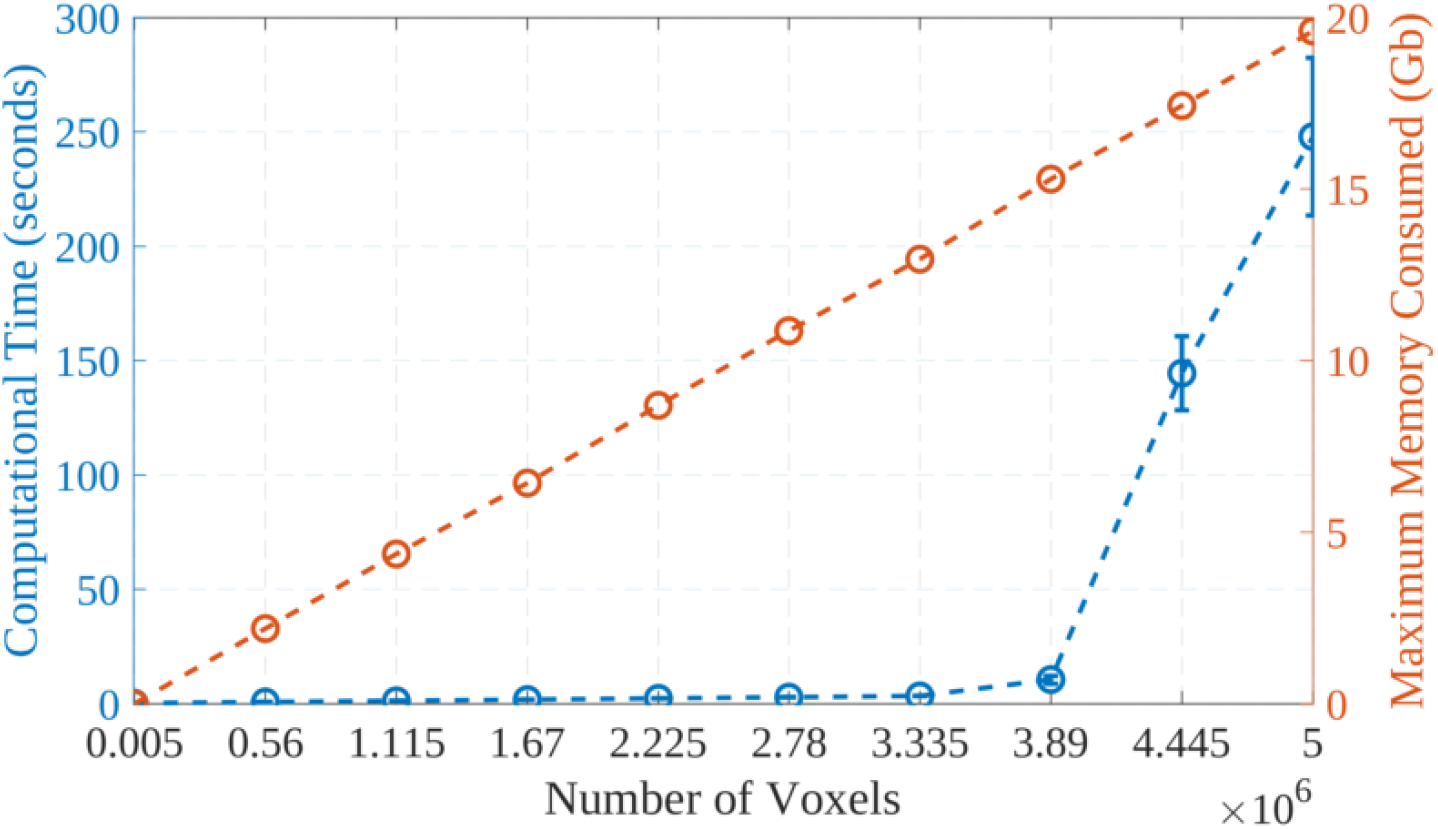
Memory and computational time requirements for the real-time pRF as a function of the number of voxels. Data points corresponding to 0s reflect < 1 Kb of memory consumed and < 0.01 seconds required for execution, respectively.

### 3.3. Poorly estimated receptive field size

At larger eccentricities our approach shows poor correspondence with the grid-search algorithm in terms of receptive field size. This is surprising giving the good correspondence between estimated and ground-truth receptive field sizes for simulated data. One potential reason for the discrepancy between our (model-free) and the grid-search approach is that the latter assumes receptive fields to have a circular shape. If receptive fields are not circular, a grid-search method may estimate receptive field sizes inaccurately. Several studies have suggested that receptive fields become increasingly elongated at higher eccentricities (Greene et al., 2014; Silson et al., 2018; Lee et al., 2013) rendering this a viable explanation for the discrepancy. An alternative explanation, assuming receptive fields are generally circular, is that the model-based grid search procedure can accurately capture sizes of receptive fields located beyond the visual field of view (the region of the visual field covered by the stimulus) whereas our model-free procedure cannot. Indeed, our model-free procedure would produce a smaller, elongated, receptive field located within the field of view if a large receptive field is located outside the field of view. Below we explore both possibilities.

#### 3.3.1. Anisotropic Model

We investigate the ability of our approach to capture elongated receptive fields by generating simulated data (similar to 2.4.1) using anisotropic Gaussians as ground-truth receptive fields:

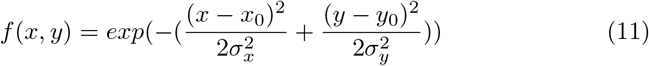

We vary *σ*_*y*_ as a ratio of *σ*_*x*_ such that the ratio between *σ*_*x*_ and *σ*_*y*_ increases with eccentricity. We first obtain *σ*_*x*_ as described in section 2.4.1. We then compute *σ*_*y*_ = *ratio ∗ σ*_*x*_; where *ratio* is *σ*_*x*_ rescaled in the range [0.5, 3]. We generate simulated 3T an 7T data with this anisotropic model with the remaining simulation parameters remaining the same as described in section 2.4.1. We define standard deviation *σ* of such anisotropic receptive fields as the geometric mean of *σ*_*x*_ and *σ*_*y*_, that is, 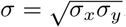. Using the geometric mean ensures that the area of an ellipse with semi-minor axis *σ*_*x*_ and semi-major axis *σ*_*y*_ is the same as a circle with radius of *σ*.

To examine whether or not our approach reliably captures the shape of the receptive fields, we visually inspect them. Figures 12 and 13 show that our approach is able to generally capture anisotropic receptive field shapes and sizes rather well. The corresponding correlation coefficients are reported in Table 7. However, as receptive fields become more elongated, our method tends to slightly underestimate their size. Interestingly, the grid-search method assuming isotropic receptive fields tends to somewhat overestimate receptive field sizes at large eccentricities. In conjunction, these effects can account for the discrepancy between the ridge-based and grid-search pRF mapping procedure. In order to analyze our approach quantitatively, we compute the JS between estimated receptive fields, ground truth receptive fields and the receptive fields obtained from grid-search (see Table 6). Note that the grid-search method yields pure Gaussians containing no anomalous activations whereas our method yields anomalous activations surrounding the receptive field. Even slight anomalies get penalized in the JS thus accounting for overall better fit observed for the grid-search method.

**Table 6:** The mean Jaccard Similarity between fast and grid-search estimated pRF parameters for **a)** 3T and **b)** 7T simulated data based on the anisotropic ground-truth receptive fields.

|  | Jaccard Similarity | baseline |
| --- | --- | --- |
| Ridge regression vs. ground truth | 0.4023 (95 % CI [0.3989,0.4057]) | 0.0567 |
| Grid search vs. ground truth | 0.6217 (95 % CI [0.6195,0.6238]) | 0.0522 |
| Ridge regression vs. grid search | 0.4207 (95 % CI [0.4171,0.4243]) | 0.0652 |
| (a) |  |  |
|  | Jaccard Similarity | baseline |
| Ridge regression vs. ground truth | 0.4470 (95 % CI [0.4439,0.4502]) | 0.0565 |
| Grid search vs. ground truth | 0.6324 (95 % CI [0.6299,0.6349]) | 0.0490 |
| Ridge regression vs. grid search | 0.4140 (95 % CI [0.4100,0.4181]) | 0.0582 |
| (b) |  |  |

**Table 7:** Correlation coefficients between fast and grid-search estimated pRF parameters for simulated **a)** 3T and **b)** 7T data based on anisotropic ground-truth receptive fields.

|  | X-coordinate | Y-coordinate | Standard Deviation |
| --- | --- | --- | --- |
| Ridge regression vs. ground truth | 0.9900 (95 % CI [0.9895, 0.9903]) | 0.9886 (95 % CI [0.9881, 0.9890]) | 0.9588 (95 % CI [0.9571, 0.9604]) |
| Grid search vs. ground truth | 0.9950 (95 % CI [0.9948, 0.9952]) | 0.9962 (95 % CI [0.9960, 0.9963]) | 0.9163 (95 % CI [0.9130, 0.9195]) |
| Ridge Regression vs. grid search | 0.9909 (95 % CI [0.9905, 0.9913]) | 0.9891 (95 % CI [0.9887, 0.9896]) | 0.9122 (95 % CI [0.9088, 0.9156]) |
| (a) |  |  |  |
|  | X-coordinate | Y-coordinate | Standard Deviation |
| Ridge regression vs. ground truth | 0.9915 (95 % CI [0.9912, 0.9919]) | 0.9940 (95 % CI [0.9938, 0.9942]) | 0.9615 (95 % CI [0.9599, 0.9630]) |
| Grid search vs. ground truth | 0.9971 (95 % CI [0.9970, 0.9972]) | 0.9972 (95 % CI [0.9971, 0.9973]) | 0.9215 (95 % CI [0.9184, 0.9245]) |
| Ridge Regression vs. grid search | 0.9923 (95 % CI [0.9920, 0.9926]) | 0.9927 (95 % CI [0.9924, 0.9930]) | 0.9115 (95 % CI [0.9081, 0.9149]) |
| (b) |  |  |  |

**Figure 12:**
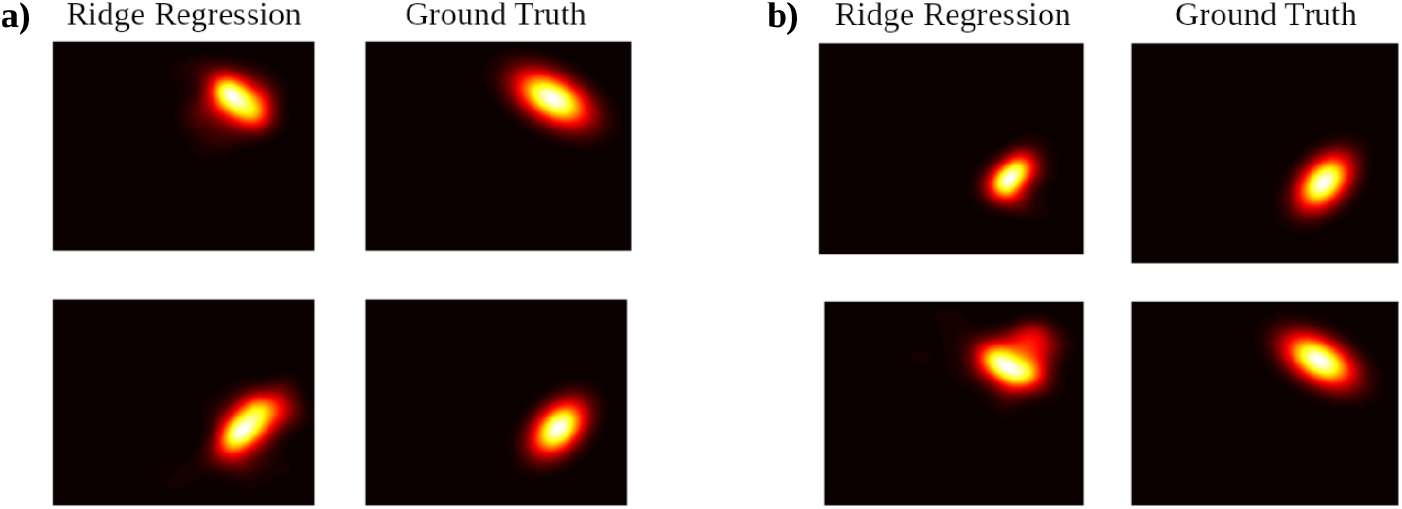
Comparison of ridge-estimated and anisotropic ground-truth receptive fields. **a)** Small (top) and large (bottom) estimated and ground-truth receptive fields for simulated 3T data. **b)** Small (top) and large (bottom) estimated and ground-truth receptive fields for simulated 7T data.

**Figure 13:**
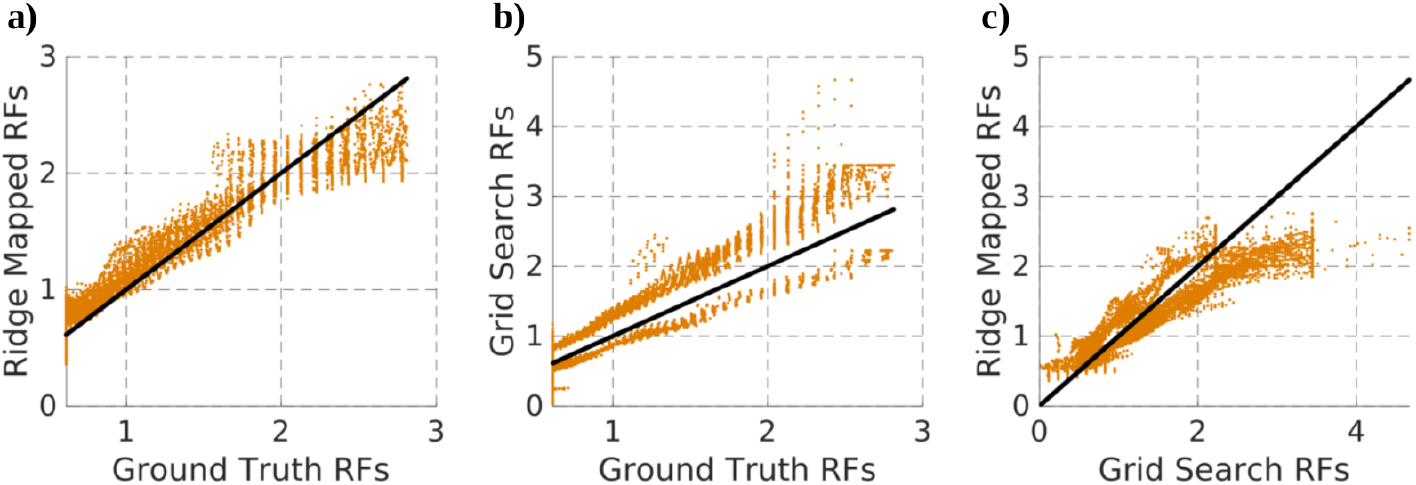
Comparison of receptive field size estimates and ground truth. **a)** Sizes estimated using our fast procedure vs ground truth sizes. **b)** Sizes estimated using grid-search vs ground truth sizes. **c)** Sizes estimated using our fast procedure vs grid-search estimates. Al results are based on simulated 3T. Results for simulated 7T data are shown in C.20

#### 3.3.2. Receptive Fields Beyond the Field of View

Next we examine to what extent our approach fails to effectively map receptive fields that (partially) lie beyond the field of view. For such receptive fields our approach maps the part that is within the field of view as an anisotropic receptive field with a relatively smaller size. The grid-search method, on the other hand, estimates these receptive fields correctly (see figure 14). This can account for the discrepancy between receptive field estimates between our method and the grid-search procedure.

**Figure 14:**
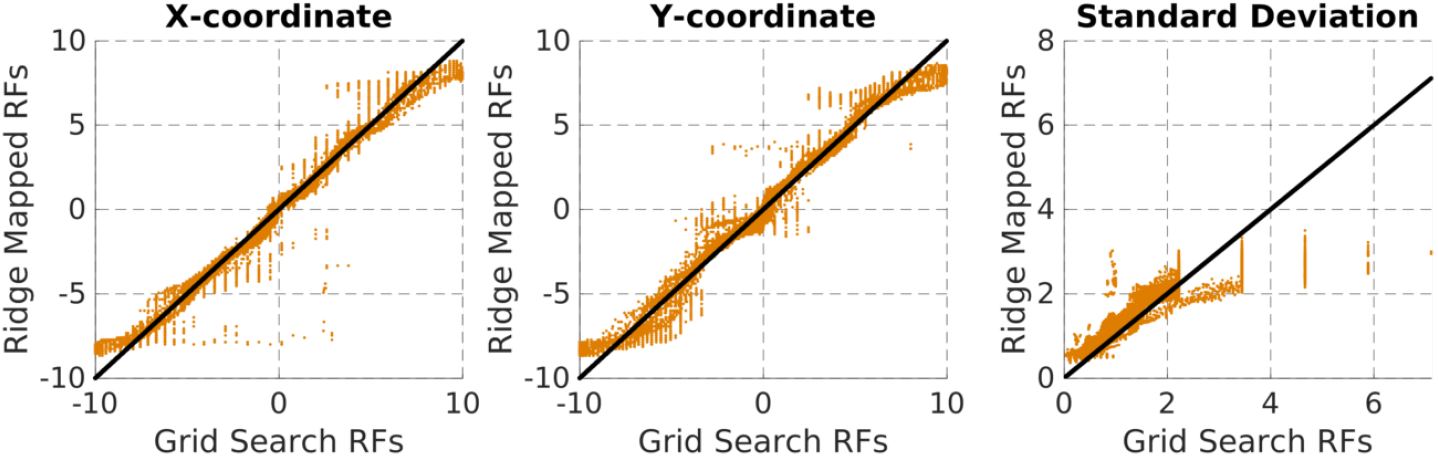
Fast procedure vs grid-search estimated pRF parameters for simulated 3T data. A line with a slope of 1 is included as a reference.

In order to map larger receptive fields, we recommend using a stimulus that covers a larger field of view. Figure 15 shows the relationship between the field of view, receptive field eccentricity and the maximum reliable estimate of receptive field size using our method. We define a metric which allows us to determine the largest estimated receptive field which is also predicted with high accuracy. We first normalize the sizes of estimated (*σ*_*r*_) and ground-truth standard deviations (*σ*_*t*_) to the range [0, 1]. Then, we select the largest reliable standard deviation as 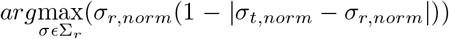. We simulate receptive fields located at a range of eccentricities [0, 30] with a range of sizes [0.5, 30] and estimate them with stimuli covering a range of field of views [5, 25]. As can be appreciated from the figure, accuracy of receptive field sizes obtained from our fast parameter estimation procedure depends on the field of view and on eccentricity. This should be taken into account when interpreting results obtained from our method.

**Figure 15:**
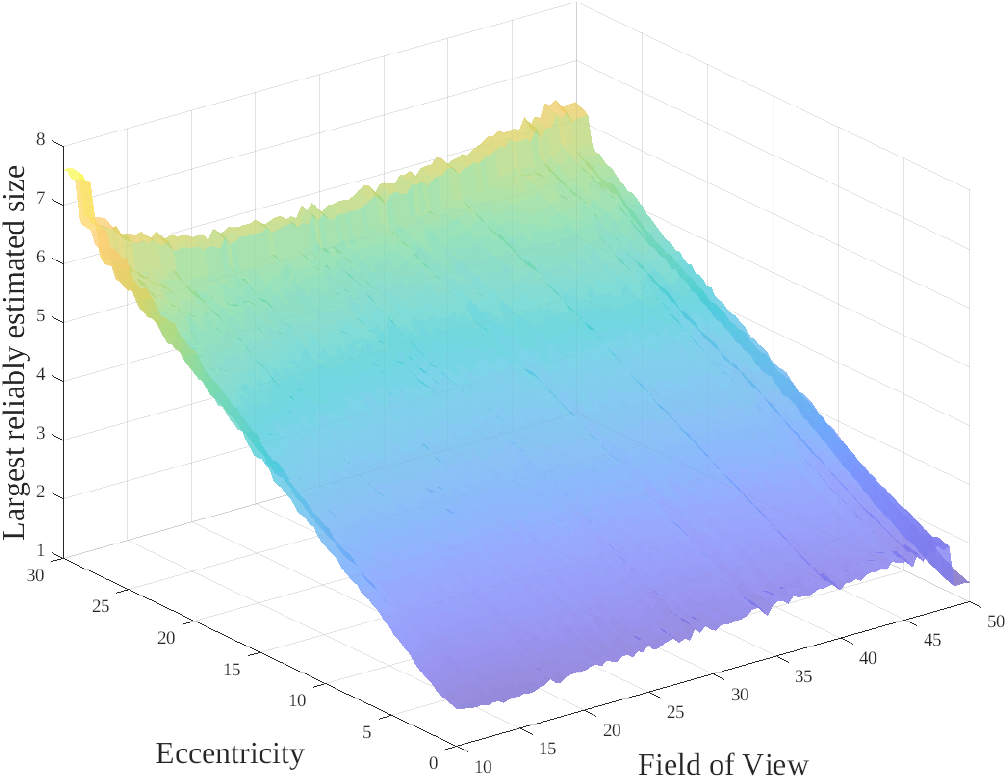
Largest reliably estimated receptive field size as a function of eccentricity and field of view. The surface plot was smoothed using *smooth2* (Hilands, 2020)

## 4. Discussion and Conclusion

We propose a fast and model-free approach for receptive field mapping and pRF parameter estimation that is suitable for real-time applications. A voxel-to-pixel map typically is a huge data matrix which requires much computer memory in order to be stored and to be operated on; rendering operations slow. To reduce data by more than 90%, we encode the stimulus using tile coding and hashing. This lowers memory requirements and hence strongly reduces computational time. We evaluated our approach on simulated as well as real empirical data in terms of computational times, fidelity of estimated receptive field shapes and parameters and the suitability of estimated pRF shapes for projecting cortical activity back into the visual field.

We find that our approach is extremely fast at mapping pRFs and estimating their parameters with computational times in the order of seconds and ~ 1 minute, respectively. Specifically, because our approach can successfully estimate receptive field shapes for large amounts of voxels in mere seconds, it is straightforward to identify the best performing (i.e. visually responsive) voxels by conducting a quick cross-validation procedure. This allows limiting parameter estimation to these voxels and thus to keep computational time low for this process as well. This also eliminates the need of using a pre-defined mask. Furthermore, cross-validation is performed in batches and we provide the option of adjusting the batch size which can further speed up parameter estimation.

In terms of fidelity, we observe excellent correspondences between estimated pRFs and ground-truth pRFs both in terms of shapes and parameters for simulated data. For empirical data, we observe excellent correspondence between pRF locations estimated from our procedure using grid-search on an isotropic Gaussian model. However, for pRF size (standard deviation) of the receptive fields results of the two methods correspond less well. In particular for larger eccentricities correspondence is poor. This is because our method finds anisotropic (elongated) receptive fields. Similar observations were also reported in (Lee et al., 2013) and it was suggested that the receptive fields tend to be anisotropic towards the edge of the stimulus space. The authors argue that when a receptive field partially lies outside the stimulus space, the part of the receptive field that lies inside may be incorrectly identified as having an oval shape. This would be in line with recent studies arguing that receptive fields are generally circular in shape (Lerma-Usabiaga et al., 2020). Other studies have suggested that receptive fields do become increasingly elongated at higher eccentricities. To investigate how both possibilities affect our mapping procedure we conducted additional, unplanned, analyses.

First, we simulated data based on elliptical receptive fields. We observe excellent correspondence between pRF parameters from our approach and ground-truth parameters. Our algorithm estimates the size of such elliptical receptive fields better than the grid-search method. This means that our method is flexible and freer in capturing the shape of the receptive fields than model-based methods. As such, our method is in principle able to capture the true shape of a receptive field. However, an analysis of how receptive field size estimates are effected by their eccentricity and the visual field of view revealed that estimates are only accurate within a certain region of the visual field of view. The flexibility of our method comes at the cost of an inability to deal with large, circular, receptive fields that lie beyond the field of view (i.e. outside the region of stimulation). This is in line with the observation that linear encoding methods (such as ridge regression) fail to reliably estimate large receptive fields (Lage-Castellanos et al., 2020); or rather the receptive fields that partially lie beyond the field of view.

From results shown figure 15 it is possible to derive up to which eccentricity receptive field size estimates are reliable given the field of view of a particular experimental setup. In order to map receptive fields outside of that region, we recommend either using a larger stimulus space or to use a model-based algorithm, such as a Levenberg-Marquadt algorithm or the grid-search method, to fit pRF parameters. To benefit from the fast procedures described here as well as the accuracy of grid-search, it is recommended to utilize the cross-validation procedure included in our method to identify visually responsive voxels and hence to reduce the total number of voxels for which grid-search needs to be performed.

Importantly, for the purpose of projecting cortical activations back into the visual field, the true shape of receptive fields at the edges of the visual field do not matter. Indeed, as can be seen from figure 10, our model-free approach faithfully reconstructs the letter shapes from their associated BOLD activity. Recognizable reconstructions of these shapes was possible even though data underwent real-time preprocessing which is generally considered being of lower quality than offline preprocessing. This highlights that our method is suitable for real-time applications such as content-based BCI letter-speller systems.

In that context it is also important to highlight that the results reported here were obtained using a single set of hyperparameters (learning rate, FWHM and shrinkage factor) except for reconstruction of mental imagery where we use higher shrinkage factor. While the choice of hyperparameters can affect mapping procedure and parameter estimation (refer to Appendix B), the set of hyperparameters used here produced robust results across participants, field strength and pre-processing procedures.

In conclusion, we present an extremely fast and flexible pRF mapping approach which can be either used in parallel with data acquisition (online gradient descent) or after the data has been fully acquired (ridge regression). This opens the door for real-time applications that rely on pRF estimates such as BCI speller systems. We also propose a fast method to estimate pRF parameters. A limitation of this method and model-free approaches in general is that receptive fields partially lying beyond the stimulus space are dealt with poorly. This can be remedied by combining fast estimation of receptive fields with a subsequent grid-search step.

## 5.Acknowledgements

This project has received funding from the European Union’s Horizon 2020 Research and Innovation Programme under Grant Agreement Numbers 945539 (HBP SGA3) and 779860 (ERC-2017-PoC).

## Appendix A. Hyperparameter sharing between gradient descent and ridge regression

Here we show that, under reasonable assumptions, the learning rate for gradient descent and the regularization parameter in ridge regression are inversely related. In a real-time setting, each iteration of online gradient descent corresponds to observing a single data point (time volume). Alternatively, one might consider performing an offline gradient descent with a single iteration where a single batch contains the entire dataset. That is, referring to equation 8, *n* = 1 and *n* − 1 = 0. If we assume that we initialize *θ*_0_ ← 0, where *θ*_0_ is the optimal solution, then we get:

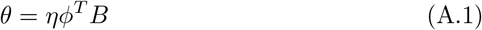

And from equtaion 5, we have:

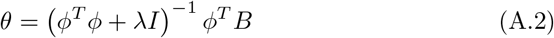

For the case that 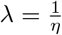 and that λ is sufficiently large such that Φ^*T*^ Φ+*λ*I ≈ λ*I*,

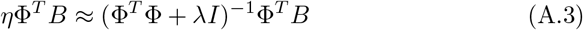

Thus, if we use 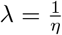 with a sufficiently large λ, ridge regression and online gradient descent yield similar results.

## Appendix B. Effect of hyperparameters on mapping procedure

The set of hyperparameters involved in the mapping procedure are learning rate (or regularization parameter), FWHM of hashed Gaussians and shrinkage. Note that we do not address learning rate and regularization parameter separately since we assume them to be the inverse of each other (refer to Appendix A). For all the experiments reported in this paper, we use the same set of hyperparameters (except for projecting cortical activity back into the visual field, which benefits from slightly higher shrinkage).

In order to understand how the choice of hyperparameters can affect mapping procedure, we fine-tune them by minimizing an objective function using Bayesian Optimization. Bayesian Optimization enables us to visualize the model mean (estimated objective function surface). For objective function or loss function we use Jaccard Distance which can be defined as:

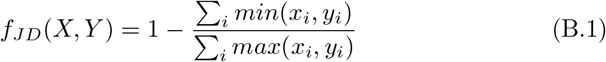

In figure B.16, we present objective function models of two cases: B.16**a** where we keep shrinkage constant and B.16**b** where keep learning rate constant. It can be seen from figure B.16**a** that the mapping procedure is not very sensitive to learning rate. However, for the relation proved in A.3 to hold true, we recommend using a small value of learning rate (< 1). Shrinkage does not have any effect on mapping itself, since it is used after the mapping procedure to remove abnormal pixels surrounding the receptive fields. Using a large value of shrinkage will reduce the size of the receptive field. FWHM, however, has a direct effect on the mapping procedure. Figure B.17 shows how a combination of FWHM and shrinkage affect the shape of the mapped receptive fields. Using a large FWHM would result in a large overlap between the stimulus and hashed Gaussians, thereby over-encoding the presence of the stimulus. As a result, the mapping procedure would overestimate the size of the receptive fields. This effect is clear from figure B.17, where we visually compare the receptive fields (for the same voxel) obtained using different values of FWHM. The shrinkage and FWHM have an opposite effect on the receptive fields. Hence it is important to use a balanced choice of FWHM and shrinkage in order to obtain optimal receptive fields.

**Figure B.16:**
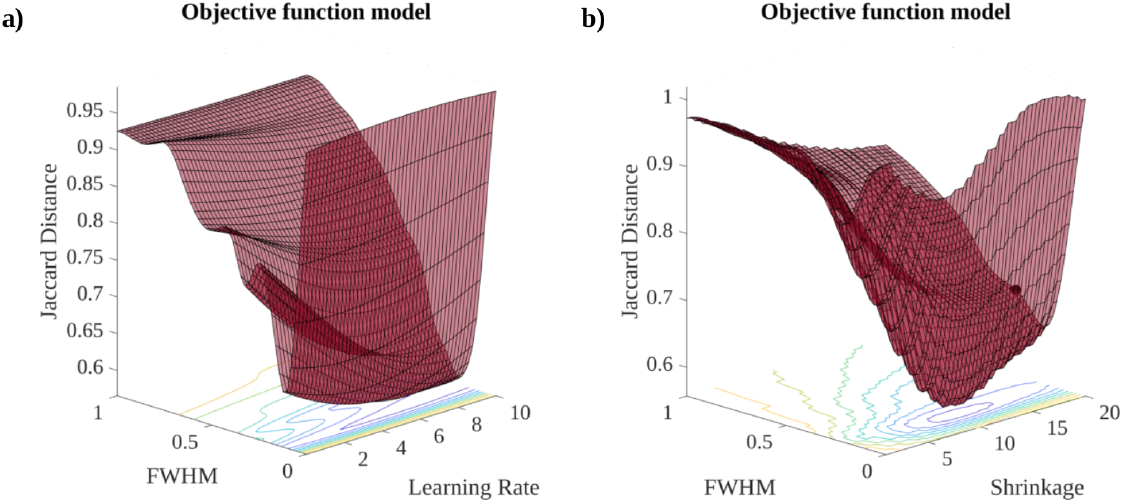
Estimated objective function model for **a)** the learning rate and the FWHM (the shrinkage was set to 6 **b)** the shrinkage and FWHM (the learning rate was set to *η* = 0.1. The hyperparameters were optimized for Jaccard Distance between mapped receptive fields and ground-truth receptive fields based on 3T-like simulated data. The optimization was performed using Bayesian Optimization. The optimization was stopped after 40 evaluations.

**Figure B.17:**
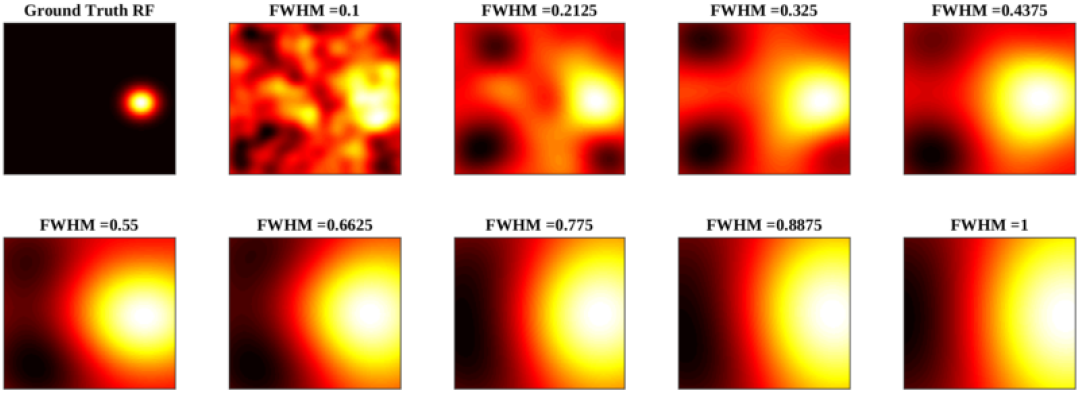
The effect of FWHM of hashed Gaussians on mapped receptive fields. The receptive field on the top left is the ground-truth receptive field based on 3T-like simulated data. The rest of the receptive fields were mapped using different FWHMs in the range [0.1, 1]. The learning rate was kept constant to 0.1 and shrinkage was not used. Note that, we use FWHM relative to resolution of stimulus space and hence it is restricted to the range [0, 1].

## Appendix C. Supplementary Figures

**Figure C.18:**
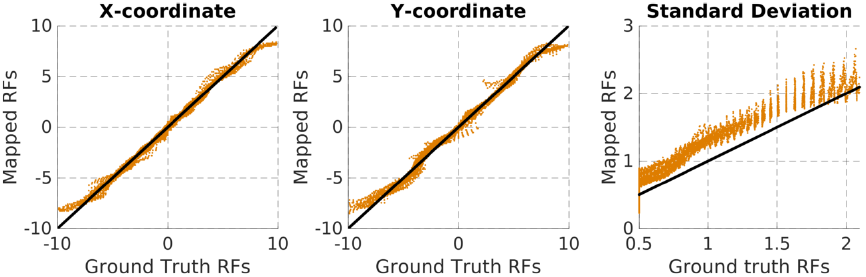
Scatter plots between the pRF parameters (location and size) estimated using the fast estimation technique and the ground-truth pRF parameters for 7Tesla-like simulated data. The voxels lying beyond the radius of measured visual field (maximum eccentricity) were ignored for estimating pRF parameters.

**Figure C.19:**
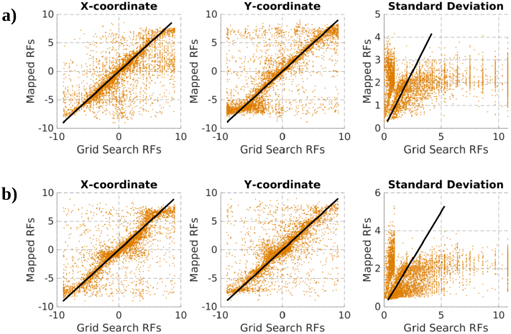
Scatter plots between the pRF parameters (location and size) estimated using fast estimation technique (ridge regression) and the grid search method on 3 Tesla empirical data. The subfigures **a** and **b** correspond to subjects 02 and subject 03 respectively.

**Figure C.20:**
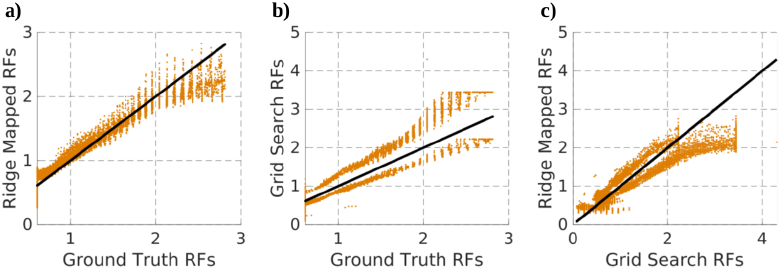
Comparison of receptive field size estimates and ground truth for 7T empirical data. **a)** Sizes estimated using our fast procedure vs ground truth sizes. **b)** Sizes estimated using grid-search vs ground truth sizes. **c)** Sizes estimated using our fast procedure vs grid-search estimates.

**Figure C.21:**
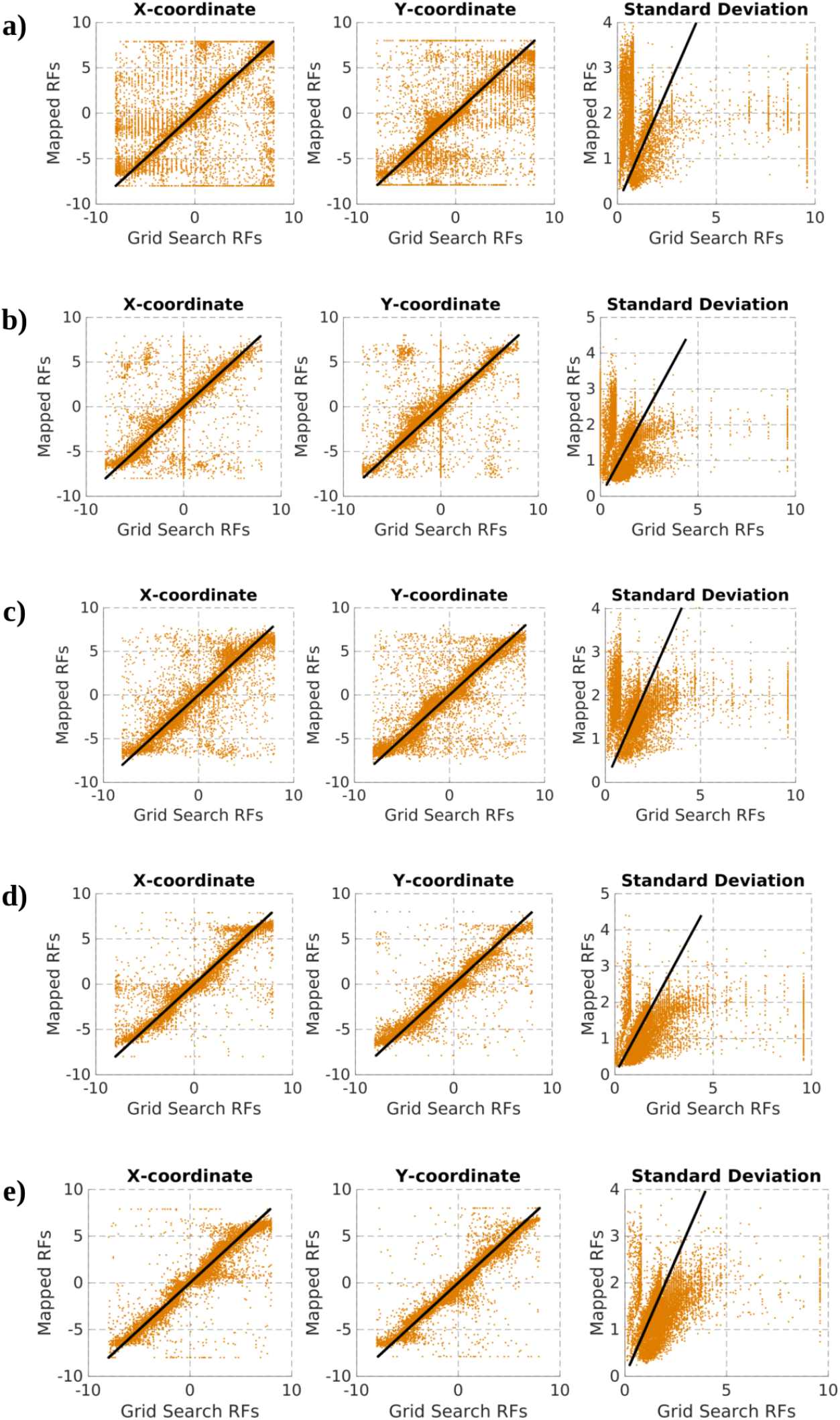
Scatter plots between the pRF parameters (location and size) estimated using fast estimation technique (ridge regression) and the grid search method on 3 Tesla empirical data. The subfigures **a**, **b**, **c**, **d**, **e** correspond to subjects 01, 02, 04, 05 and 06 respectively.

## Notes

https://github.com/ccnmaastricht/real_time_pRF

### Competing Interest Statement

The authors have declared no competing interest.

